# Metabolic Engineering Reveals *LUP5* as a Determinant of Saponin Composition and Insect Resistance in *Barbarea vulgaris*

**DOI:** 10.1101/2025.08.04.665341

**Authors:** Jincheng Shen, Jan Günther, Sebastian Kjeldgaard-Nintemann, Pablo D. Cárdenas, Søren Bak

## Abstract

Plant–insect coevolution has been a major driver of specialized metabolite diversification, yet the genetic basis of natural variation in defensive chemistry remains poorly understood. The wild crucifer *Barbarea vulgaris* comprises two ecotypes, an insect-resistant G-type and a susceptible P-type, characterized by distinct triterpenoid saponin profiles. To investigate the causal relationship between saponin composition and insect resistance, we established a stable transformation system for *B. vulgaris*. Expression of the G-type β-amyrin synthase gene *LUP5* in the susceptible P-type conferred up to a 95% reduction in *Plutella xylostella* feeding, accompanied by increased accumulation of three hederagenin-derived monodesmosidic saponins. Comparison of *LUP5* expression driven by its native promoter and by the constitutive 35S promoter revealed that the native promoter leads to increased hederagenin accumulation and is activated later in development, which may prevent early metabolic stress and allow coordinated expression of downstream pathway genes. In contrast, silencing of *CYP72A552* by RNAi decreased total hederagenin levels by approximately 40% without affecting resistance, indicating threshold-dependent defense. Our results provide direct *in planta* evidence that *LUP5* is a key determinant of natural variation in insect resistance in *B. vulgaris*, underscoring the pivotal role of the saponin backbone in herbivore deterrence. By linking promoter activity to metabolite structural diversity, this work provides mechanistic and conceptual insight into how plants coordinate specialized metabolism and defense.

## Introduction

Plants have evolved a wide array of specialized metabolites to deter insect herbivores (Fürstenberg-Hägg, Zagrobelny and Bak, 2013; Yactayo-Chang et al., 2020; Beran and Petschenka, 2022). Among these, triterpenoid saponins, which consist of a hydrophobic sapogenin backbone and one or more hydrophilic sugar moieties, represent a structurally diverse and ecologically important class of defensive compounds. Due to their structural similarity to sterols and steroid hormones, triterpenoid saponins have been hypothesized to interact with eukaryotic cell membranes (Cárdenas, Almeida and Bak, 2019). In addition, their inherent bitterness may deter herbivores, contributing to their role as effective defense agents (Augustin et al., 2011). Thousands of triterpenoid saponins have been reported to date (Xu, Fazio and Matsuda, 2004; Augustin et al., 2011; Moses, Papadopoulou and Osbourn, 2014; Huang et al., 2020). Importantly, specific saponin structures have been implicated in insect resistance, highlighting the functional significance of their chemical diversity (Stevenson, Nyirenda and Veitch, 2010; Cui et al., 2019; Hussain et al., 2019; Liu et al., 2019; Dolma et al., 2021).

The structure–activity relationships underlying saponin bioactivity are critical to their defensive function. For example, hederagenin 3-*O*-monoglucoside has higher toxicity and antifeedant activity than oleanolic acid 3-*O*-monoglucoside (which lacks C-23 hydroxylation), and gypsogenic acid 3-*O*-monoglucoside (which is further oxidized at C-23 to the ketone) (Liu et al., 2019). Additionally, monodesmosidic hederagenin and oleanolic acid exhibited higher toxicity and antifeedant activity than their bidesmosidic counterparts (Tian et al., 2021; Dervishi et al., 2025, in press). These observations suggest that even subtle modifications to the sapogenin backbone, glycosylation pattern, or sugar linkage can significantly alter saponin bioactivity. This structural diversity makes saponins a valuable system for studying molecular evolutionary mechanisms of specialized metabolism, particularly the structure– activity relationships underlying their ecological functions in insect defense. Despite growing knowledge of these structure–activity relationships, the genetic and biochemical mechanisms underlying saponin diversification remain poorly understood in most species. One particularly promising ecological model to explore this question is the wild crucifer *Barbarea vulgaris*, which naturally segregates in two ecotypes or chemotypes that alter in both saponin profiles and insect resistance.

*B. vulgaris* has emerged as a powerful ecological model for investigating how specialized metabolites contribute to herbivore resistance (Kuzina et al., 2011; Hauser, Toneatto and Nielsen, 2012; Byrne et al., 2017). The species comprises two ecotypes: the glabrous insect-resistant G-type, and the pubescent insect-susceptible P-type (Agerbirk, Olsen and Nielsen, 2001). This intraspecific divergence likely arose from isolation in different Ice Age refugia (Hauser, Toneatto and Nielsen, 2012) and is associated with marked differences in saponin composition (Kuzina et al., 2009; Khakimov et al., 2012; Khakimov et al., 2016). Gene duplication events have further contributed to the chemical divergence and insect resistance (Khakimov et al., 2015; Liu et al., 2019). Insect resistance in G-type *B. vulgaris* correlates with four oleanane-type defensive saponins, including oleanolic acid cellobioside, hederagenin cellobioside, gypsogenin cellobioside, and 4-epihederagenin cellobioside (Agerbirk et al., 2003; Kuzina et al., 2011). Among these, hederagenin cellobioside is particularly toxic and confers G-type insect resistance (Agerbirk et al., 2003; Nielsen et al., 2010; Kuzina et al., 2011).

The biosynthetic pathway leading to hederagenin cellobioside in G-type *B. vulgaris* has been elucidated up to the first glycosylation step (Fig. 1). The oxidosqualene cyclase *LUP5* is highly expressed in the G-type and primarily produces β-amyrin (Khakimov et al., 2015). Subsequent enzymatic steps involve CYP716A80, CYP72A552, and UGT73C11, which sequentially convert β-amyrin into oleanolic acid, hederagenin, and hederagenin 3-*O*-glucoside (Fig. 1) (Augustin et al., 2012; Khakimov et al., 2015; Liu et al., 2019). The enzyme responsible for the final step in forming hederagenin cellobioside remains unidentified (Erthmann, Agerbirk and Bak, 2018). In contrast, *LUP5* is expressed at very low levels in *B. vulgaris* P-type and primarily produces α-amyrin due to two amino acid substitutions as compared to the *LUP5* in the G-type (Günther et al., 2021). Despite 98% sequence identity at the amino acid level, the G- and P-type LUP5 enzymes differ markedly in both their gene expression and product profiles. The P-type plants preferentially express *LUP2,* an oxidosqualene cyclase over 80% identical at the amino acid level to *LUP5*, but it produces predominantly lupeol (98%) and only 2% β-amyrin-derived saponins (Khakimov et al., 2015). Notably, the genes encoding for the remaining enzymes responsible for the biosynthesis of hederagenin 3-*O*-glucoside (CYP716A80, CYP72A552, and UGT73C11) are present in both P-type and G-type and exhibit similar substrate and product specificity (Augustin et al., 2012; Khakimov et al., 2015; Liu et al., 2019). However, the *cyclases* are differentially regulated suggesting that upstream regulation and oxidosqualene cyclase product specificity may be key drivers of saponin structural and insect resistance difference. Additionally, CYP72A552, which catalyzes the C23 hydroxylation of oleanolic acid to hederagenin, the precursor of the more toxic hederagenin-type saponins, has not yet been validated *in planta* for its ecological contribution to insect resistance in *B. vulgaris*.

**Fig. 1.**
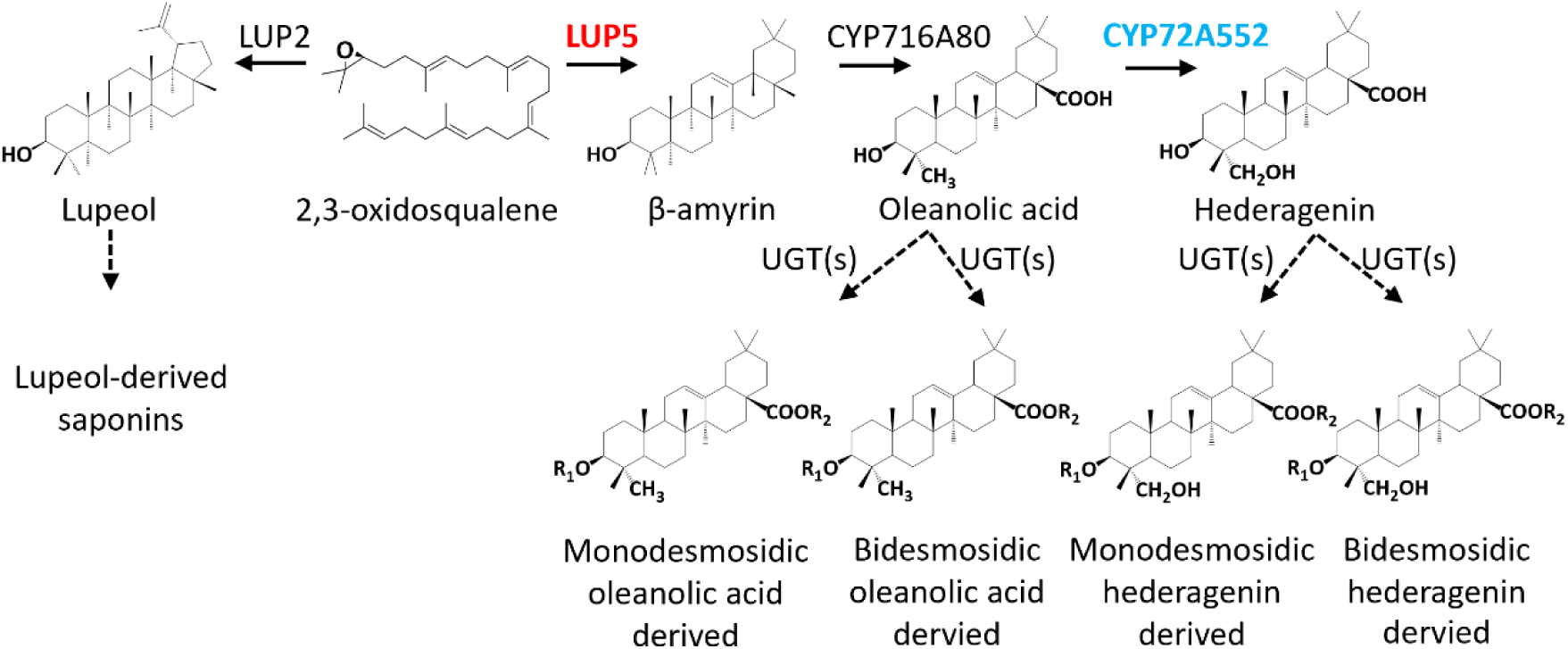
Biosynthetic pathways for lupeol and β–amyrin derived triterpenoid saponins in *B. vulgaris*. Monodesmosidic saponin, a single sugar chain attached at preferentially C-3 (R1) or C-28 (R2); bidesmosidic saponin, sugar chains attached at both C-3 (R1) and C-28 (R2).

To investigate the genetic and biochemical basis of saponin-mediated insect resistance, we leveraged the natural variation in *B. vulgaris* to develop a model system linking *in planta* metabolic engineering of specialized metabolism with ecological function. We provide functional evidence that engineering saponin biosynthesis through manipulation of *LUP5* expression substantially enhances insect resistance in this wild crucifer and explains the molecular basis of the difference in insect resistance between the G- and P-type *B. vulgaris*. By comparing constructs driven by the native *LUP5* and constitutive 35S promoters, we further reveal how promoter activity shapes metabolite structural diversity. These findings offer new mechanistic insight into plant–insect coevolution and established a model system for dissecting genotype– phenotype relationships in plant chemical specialization.

## Materials and methods

### Plant and insect materials

Seeds of *Barbarea vulgaris* G-type (glabrous, accession B44, Herbarium-code: CP0057358) and P-type (pubescent, accession B4, herbarium-code: CP0057347) were provided by Associate professor Niels Agerbirk *(Agerbirk et al., 2021)* and used for stable transformation. Both transformed and wild-type plants were cultivated in soil (Krukvaxtjord med lera och kisel, SW Horto AB, Hammenhog, Sweden) in a greenhouse maintained at 19 °C under a 16 h light/8 h dark photoperiod. Irradiance was supplemented with LED lamps whenever natural light intensity dropped below 250 µmol m^-2^ s^-1^, with full-spectrum sunlight available when curtains were open. Plants were fertigated on Thursdays and Fridays with a nutrient solution (electrical conductivity 2.2 mS cm^-1^; pH 5.8) prepared by dissolving 100 g L^-1^ of a compound fertilizer (Pioner Basis 13-2-23+3+ME; containing 13% N, 2% P_₂_O_₅_, 23% K_₂_O, 3% MgO, plus micronutrients) together with 0.5 g L^-1^ of iron chelate (Pioner Fe-EDDHA 6%; 6% Fe) in water. *Brassica napus* plants were grown in the same greenhouse under identical conditions. *Plutella xylostella* (diamondback moth) eggs were obtained from Dr. Patrick Hughes (Boyce Thompson Institute, Ithaca, NY, USA). The laboratory colony was originally established at the New York State Agricultural Experiment Station (Geneva, NY, USA) in 1994. The insects were reared on *B. napus* at 20 °C under a 16 h light/8 h dark photoperiod in cages. Third instar *P. xylostella* larvae were used for the insect feeding assay.

### Plasmids construction

For stable transformation, the pJCV51 vector was employed to introduce the G-type *LUP5* coding sequence and to assess tissue-specific promoter activity (native *LUP5* and constitutive 35S), while pK7FWG2 was utilized for *CYP72A552* silencing (Karimi, Inzé and Depicker, 2002). The *LUP5* coding sequence, its native promoter (1988 bp), and the *CYP72A552* coding sequence were PCR-amplified from *B. vulgaris* G-type cDNA and genomic DNA (Supplementary Table S1 and S2). G-type *LUP5* expression was driven either by the 35S promoter or its native promoter, while *CYP72A552* was driven by the 35S promoter.

The pJCV51-p35S::LUP5, pJCV51-p35S::eGFP, and pK7FWG2-CYP72A552 RNAi plasmids were assembled using Gateway technology (Gateway™ BP Clonase™ II Enzyme mix, Thermo Fisher Scientific, 11789020; Gateway™ LR Clonase™ II Enzyme mix, Thermo Fisher Scientific, 11791020) (Supplementary Table S1). pDONR207 was used as the entry clone vector for both constructs (Thermo Fisher Scientific, 117207-021). Subsequently, the pJCV51-pLUP5::LUP5 and pJCV51-pLUP5::eGFP plasmids were generated by replacing the 35S promoter in pJCV51-p35S::LUP5 or pJCV51-p35S::eGFP with the G-type *LUP5* promoter using the SalI sites. Finally, the plasmids were introduced into *Agrobacterium tumefaciens* strain AGL1 via electroporation (Hayta et al., 2021).

### Stable transformation in wild crucifer *B. vulgaris*

Seed sterilization was achieved by immersing seeds in 70% ethanol for 30 s, followed by a 12.5 min treatment in 3% sodium hypochlorite (Thermo Fisher Scientific) with 0.1% Tween20 (Bio-RAD, 1706531) and thorough rinsing with sterile water. Seeds were sown on germination medium in Magenta boxes and incubated in darkness for 6–8 days (Supplementary Table S3).

Hypocotyl explants (∼3 mm) from 6–10-day-old seedlings were incubated for 15 min with *A. tumefaciens* strains (OD₆₀₀ = 0.5 in 10 mM MgCl₂, 100 µM acetosyringone) with gentle agitation and then co-cultivated in darkness for three days (Supplementary Table S3). The pJCV51-pLUP5::LUP5 and pJCV51-35S::LUP5 constructs were transformed into both *B. vulgaris* ecotypes, while pJCV51-p35S::eGFP, pJCV51-pLUP5::eGFP, and pK7FWG2-CYP72A552 RNAi were introduced only into the G-type.

After rinsing in Milli-Q water containing 1 µL/mL timentin, the explants were transferred sequentially to callus induction (one week), shoot induction (two rounds, two weeks each), and root induction media (two rounds, two weeks each) at 25°C under a 16:8 h light–dark cycle (Supplementary Table S3). Throughout these stages, the explants were maintained in a climate chamber at 25°C under a 16:8 h light-dark cycle. Kanamycin (50 ng/mL) was used for selecting transformed explants, whereas wild-type explants were cultured without kanamycin supplementation in the medium.

CYP72A552 RNAi plants were confirmed by NPTII (neomycin phosphotransferase II) ELISA Kit (Agdia, PSP 73000/0288). All the other transgenic plants were confirmed by RFP fluorescence (for *LUP5* expression and promoter activity constructs). Fluorescence was analyzed on a Leica M205FA fluorescence dissection microscope (Leica Microsystems). RFP and eGFP were imaged using the dsRed plant (excitation 546/10 nm; emission 600/40 nm) or ET GFP filters (excitation 470/40 nm; emission 525/50 nm), respectively. Subsequently, the confirmed plants were transferred to soil and grown in a greenhouse at 19°C under a 16:8 h light–dark cycle.

### Choice insect feeding assay

Insect resistance was evaluated using at least three individual plants (biological replicates) per transgenic line. For each transgenic plant, five transgenic and five wild-type leaf discs (1.57 cm² each) were alternately arranged in a petri dish (9.4 x 1.6 cm) (Greiner BIO-ONE, 633180) with filter paper moistened with 2 mL of water. After a 3– 5-hour starvation period, ten third-instar *P. xylostella* larvae were placed at the center of the plate, following a modified protocol from Liu et al (Liu et al., 2019). Leaf consumption was quantified when half of the wild-type leaves were consumed using ImageJ (approx 7.5 h).

### Preparation of plant extracts and metabolite analysis

The extraction of saponins and sapogenins was adapted from Khakimov et al. (2016). *B. vulgaris* leaf samples were ground with liquid nitrogen using a mortar and pestle. For saponin detection, 100 mg of leaf powder was weighed into 1.5 mL Eppendorf tubes and extracted with 300 µL of 85% methanol (v/v) via sonication for 30 minutes, followed by centrifugation at 16,000 *×* g for 10 minutes. Supernatants (200 µL) were filtered through a 0.22-μm filter before LC-qTOF-ESI-MS/MS analysis.

To detect total and free sapogenins, 200 mg of leaf powder was weighed into 2 mL Eppendorf tubes and extracted with 600 µL of 85% methanol (v/v) at 100°C while mixing at 1,400 × rpm in a thermomixer (Eppendorf, Denmark). The samples were cooled on ice and centrifuged at 16,000 × *g* for 3 min. After filtration through a 0.22- μm filter, 100 µL of supernatant was collected for free sapogenin detection via LC-qTOF-APCI-MS/MS. An additional 300 µL of supernatant was collected and dried under nitrogen flow. To cleave off sugar residues, samples were treated with 500 µL of 2 M hydrochloric acid at 100°C while mixing at 1,400 × rpm for 1.5 hours in a thermomixer (Eppendorf, Denmark). After cooling on ice, a double volume of ethyl acetate was added to the acid-water mixture, followed by vortexing and centrifugation at 3,500 × *g* for 5 minutes. The samples were extracted three times with ethyl acetate to collect sapogenins. To remove residual acid, an equal volume of Milli-Q water was added, the samples were subsequently vortexed and centrifugated at 3,500 × *g* for 5 minutes. This washing step was repeated three times. The final extracts were dried under nitrogen flow, re-solubilized in 300 µL of 100% methanol, and filtered through a 0.22- μm filter before total sapogenin detection via LC-qTOF-APCI-MS/MS.

LC-qTOF-ESI-MS/MS was performed according to Liu et al. (2019). LC-qTOF-APCI-MS/MS was performed as described by Patel et al. (2020). Untargeted metabolite data were analyzed using XCMS (Tautenhahn et al., 2012), and targeted saponin and sapogenin analyses were conducted with DataAnalysis software (v.4.3; Bruker, Bremen, Germany) following Khakimov et al. (2016). Hederagenin cellobioside and oleanolic acid cellobioside standards were used for saponin identification. Lupeol, β- amyrin, oleanolic acid, and hederagenin standards were employed for sapogenin identification.

### Gene expression analysis

Total RNA was extracted using the Spectrum™ Plant Total RNA Kit (Sigma-Aldrich, STRN50-1KT) and then synthesized to cDNA by iScript™ cDNA Synthesis Kit (Bio-Rad, 1708891). Relative expression of *LUP5* was quantified via qPCR using kapa SYBR® fast kit (Merck-Sigma, KK4607) (Khakimov et al., 2015). *Tubulin* (GenBank accession no. EU555191) was used as reference gene, and *LUP5* expression levels were normalized to the wild-type P-type control (Liu TongJin et al., 2016). Primer sequences are provided in Supplementary Table S1 (Khakimov et al., 2015).

## Results

### Establishment of a transformation and regeneration system in the wild crucifer *B. vulgaris*

To enable functional studies of insect resistance in *B. vulgaris* as an ecological model system, we developed a stable transformation and regeneration method for both the G-type and P-type ecotypes (Fig.1, Supplementary Fig. S1, Supplementary Table S3). To metabolically engineer the saponin compositions, we introduced the G-type *LUP5* coding sequence into both ecotypes and attempted to silence *CYP72A552* in the G-type. *LUP5* expression was driven either by the constitutive 35S promoter or the native *LUP5* promoter. Transformation efficiency, calculated as the percentage of positive transgenic plants to all obtained plants from tissue culture, reached up to 94% across experiments (Table 1). The native *LUP5* promoter exhibited approximately two-fold higher transformation efficiency than the 35S promoter in both the G- and P-type, indicating a selection pressure against *LUP5* overexpression from the 35S promoter (Table 1).

**Table 1.** A stable transformation system in *B. vulgaris* enabled up to 94% efficiency.

| Construct for transformation | Ecotype | Total numbers of plants | Positive/total |
| --- | --- | --- | --- |
| pJCV51-pLUP5::LUP5 | G-Type | 31 | 94% |
|  | P-Type | 35 | 60% |
| pJCV51-p35S::LUP5 | G-Type | 52 | 40% |
|  | P-Type | 66 | 26% |
| pJCV51-pLUP5::eGFP | G-Type | 24 | 58% |
| pJCV51-p35S::eGFP | G-Type | 20 | 55% |
| pK7FWG2-CYP72A552 RNAi | G-Type | 46 | 28% |

### Stable expression of G-type *LUP5* in the P-type confers insect resistance

To evaluate the impact of G-type *LUP5* expression on insect resistance, a choice insect feeding assay was conducted by presenting alternating wildtype and transgenic *B. vulgaris* leaf discs to diamondback moth larvae. Leaf area consumed was quantified when approximately half of the total leaf material of the wild type had been eaten (approx. 7.5 hr). The best performing *LUP5*-transformed P-type plants exhibited a significant reduction in larval consumption, ranging from 76 - 95% (Fig. 2, A and B). QPCR on the best performing 35S and *LUP5* promoter lines, confirmed that the G-type *LUP5* was transformed and expressed in the P-type both under the 35S promoter or the native G-type *LUP5* promoter (Fig. 2C). Interestingly, the 35S promoter expressed the *LUP5* at approximately sixteen-fold higher levels than that in the best pLUP5::LUP5 transformed P-type and the wildtype P- and G-type (Fig. 2C).

**Fig. 2.**
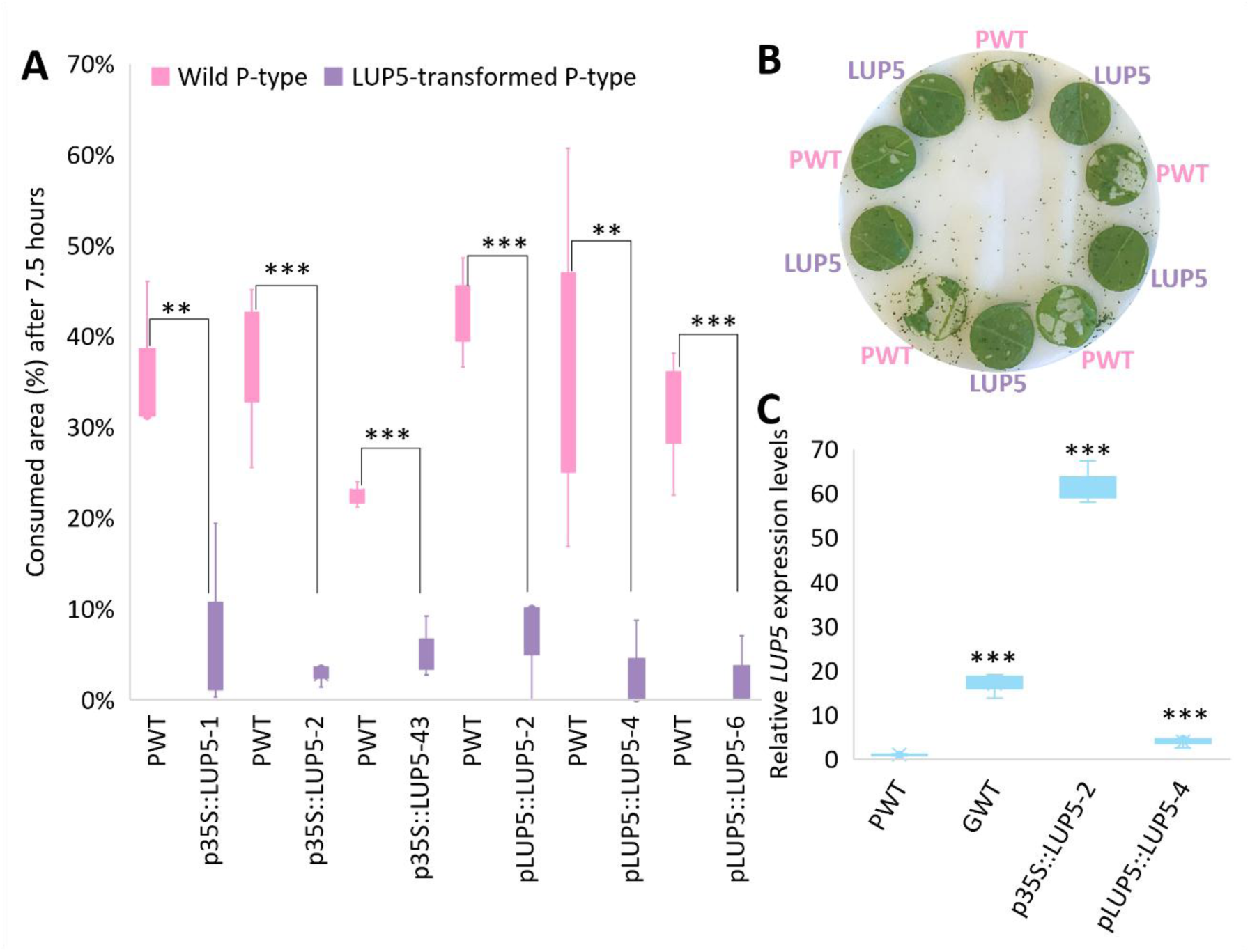
Expression of the G-type *LUP5* in *B. vulgaris* P-type reduces feeding by diamondback moth larvae. A) Consumption area (%) after 7.5 h (choice test, three individual plants per transgenic line; each replicate represents one Petri dish containing five transgenic and five wildtype leaf discs). B) Example of the insect choice feeding assay, and the resulting leaf condition after 7.5 hr insect exposure. C) Relative *LUP5* expression levels (three individual plants per transgenic line). PWT, wildtype P-type *B. vulgaris*; GWT, wildtype G-type *B. vulgaris*; p35S::LUP5, p35S::LUP5 transformed P-type *B. vulgaris*; pLUP5::LUP5, pLUP5::LUP5 transformed P-type *B. vulgaris*, LUP5, P-type *B. vulgaris* expressing the G-type *LUP5*. Statistical significance was assessed using Student’s *t*-test: *(p<0.5), ** (p < 0.05) and *** (p < 0.005).

### Expression of G-type *LUP5* induced β-amyrin-derived saponins in P-type *B. vulgaris*

Based on the highest reduction of leaf area consumed, three transgenic lines were selected for each promoter type (35S and native *LUP5*) and subjected to untargeted and targeted LCMS analyses. The untargeted metabolite analysis revealed approximately 9,500 distinct *m/z* features, each defined as a unique *m/z* and retention time (RT) pair. Features differing between *LUP5*-expressing and wild-type plants (P < 0.5) were ranked by fold change, and the top 100 increased and 100 decreased features were analyzed further. However, extraction of the *m/z* features using DataAnalysis software revealed that the majority lacked clear MS² fragmentation patterns, preventing structural annotation. Many of these features likely represent in-source fragments, background ions, or uncharacterized metabolites with inconclusive fragmentation.

Consequently, a targeted saponin analysis was conducted using DataAnalysis. *B. vulgaris* saponins were detected by extracted ion chromatograms of our previously tentatively identified aglycones and their structures were inferred from characteristic fragmentation patterns. Based on 455 m/z, 469 *m/z,* and 471 m/z, the characteristic fragmentation patterns of oleanolic acid, gypsogenin and hederagenin were tentatively identified, respectively. This targeted approach identified ten putative distinct saponins in P-type lines and thirty in G-type lines (Supplementary Table S4 and S5). All detected saponins were classified as monodesmosidic, based on their fragmentation patterns. Saponin abundance was subsequently quantified by calculating peak areas from extracted ion chromatograms (Supplementary Table S4 and S5).

The LCMS analysis revealed three hederagenin-derived monodesmosidic saponins (two hexoses; RT 10.7, 11.5, and 12.3 min) and one oleanolic acid monoglucoside (one hexose; RT 9.5 min) that accumulated at significantly higher levels in *LUP5*-transformed P-type compared to wild-type P-type (Fig. 3, A and B). The most abundant of the three hederagenin-derived saponins eluted at 11.5 min and was increased ∼four-fold in the P-type expressing *LUP5* under the native *LUP5* promoter compared to the wildtype and was ∼two-fold more abundant than in P-type plants transformed with *LUP5* under the 35S promoter (Fig. 3A). Hederagenin cellobioside (eluted at 12.3 min) was also increased ∼four-fold but remained at lower levels in pLUP5::LUP5 transformed P-type (Fig. 3A). These three hederagenin-derived saponins were also present in the control G-type plants, with hederagenin cellobioside (eluted at 12.3 min) being the most abundant hederagenin-derived saponin (Supplementary Table S5 and S8). The increased oleanolic acid monoglucoside (eluted at 9.5 min) was not detected in G-type (Supplementary Table S5). The other detected saponins remained unchanged (Supplementary Table S6).

**Fig. 3.**
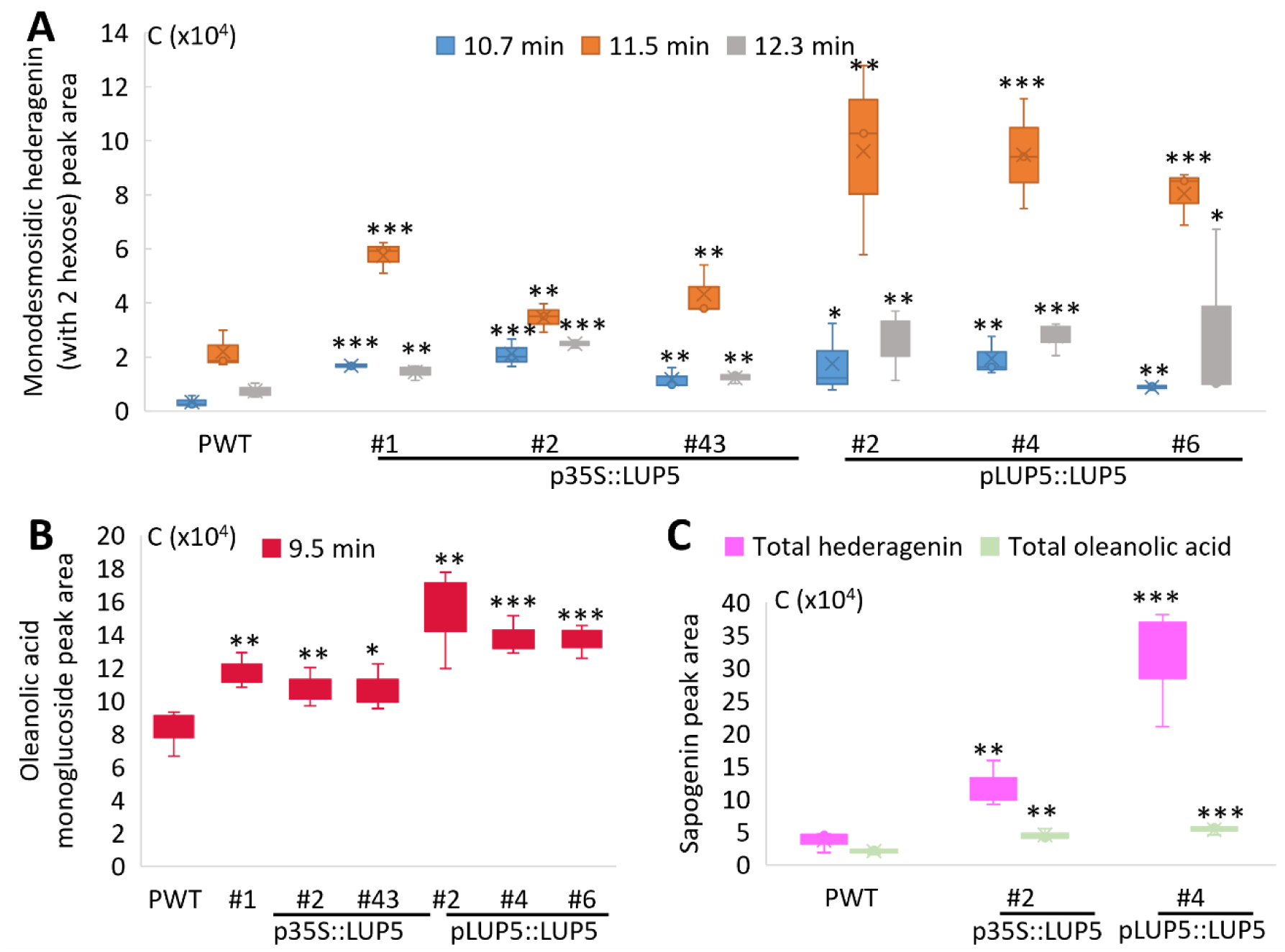
Expression of the G-type *LUP5* increased glycosylated oleanolic acid and hederagenin levels in the transgenic P-type *B. vulgaris*. A) Monodesmosidic hederagenin (with 2 hexoses) peak area (EIC: *m/z* = 841.4591 ± 0.2). B) Oleanolic acid monoglucoside peak area (EIC: *m/z* = 617.4059 ± 0.2). C) Peak area of sapogenins, including total hederagenin (EIC: *m/z* = 437.3425 ± 0.2) and total oleanolic acid (EIC: *m/z* = 439.3593± 0.2). PWT, wildtype P-type *B. vulgaris*; p35S::LUP5, p35S::LUP5 transformed P-type *B. vulgaris*; pLUP5::LUP5, pLUP5::LUP5 transformed P-type *B. vulgaris*. Error bars represent the standard deviation of the mean from three individual plants. Statistical significance was assessed using Student’s *t*-test: *(p<0.5), ** (p < 0.05) and *** (p < 0.005).

To evaluate the effect of G-type *LUP5* expression on P-type *B. vulgaris* saponin aglycone biosynthesis and accumulation, levels of free and total sapogenins were analyzed. Free sapogenins were measured from untreated samples, representing aglycones that occur naturally in a non-glycosylated form. Total sapogenins were defined as the sum of free and sugar-bound sapogenins, the latter released through hydrochloric acid mediated hydrolysis of glycosylated saponins. A transgenic line (with three vegetative clones) from both p35S::*LUP5* and pLUP5::*LUP5*-transformed P-type *B. vulgaris* was selected based on reduced leaf area consumption and increased accumulation of four saponins (Fig. 2A, Fig. 3, A and B). This analysis revealed a significant increase in total hederagenin and oleanolic acid levels in *LUP5*-transformed P-type plants (Fig. 3C), while total lupeol, β-amyrin, and free sapogenins remained undetectable (Supplementary Table S7). Higher accumulation of total hederagenin was observed in plants expressing *LUP5* under the native promoter compared to those under the 35S promoter (Fig. 3C).

To investigate how promoter choice affects gene expression and metabolic composition, we compared the previously described transgenic P-type lines expressing *LUP5* under either the constitutive 35S or the native *LUP5* promoter. Interestingly, although the 35S promoter drove much higher *LUP5* expression levels compared to the native promoter (Fig. 2C), this did not result in proportionally increased accumulation of β-amyrin–derived saponins (Fig. 3A–C). To investigate this, we made eGFP lines with the *LUP5* and 35S promoter respectively. Analysis of these lines the 35S promoter was already active at the shoot induction stage and consistently drove stronger fluorescence signals, whereas the *LUP5* promoter only became active only about four weeks after transfer to soil (Fig. 4A-F). These observations indicate that the two promoters differ mainly in temporal activation and relative expression strength, which likely contributes to the observed variation in transcript levels, metabolite accumulation, and transformation efficiency.

**Fig. 4.**
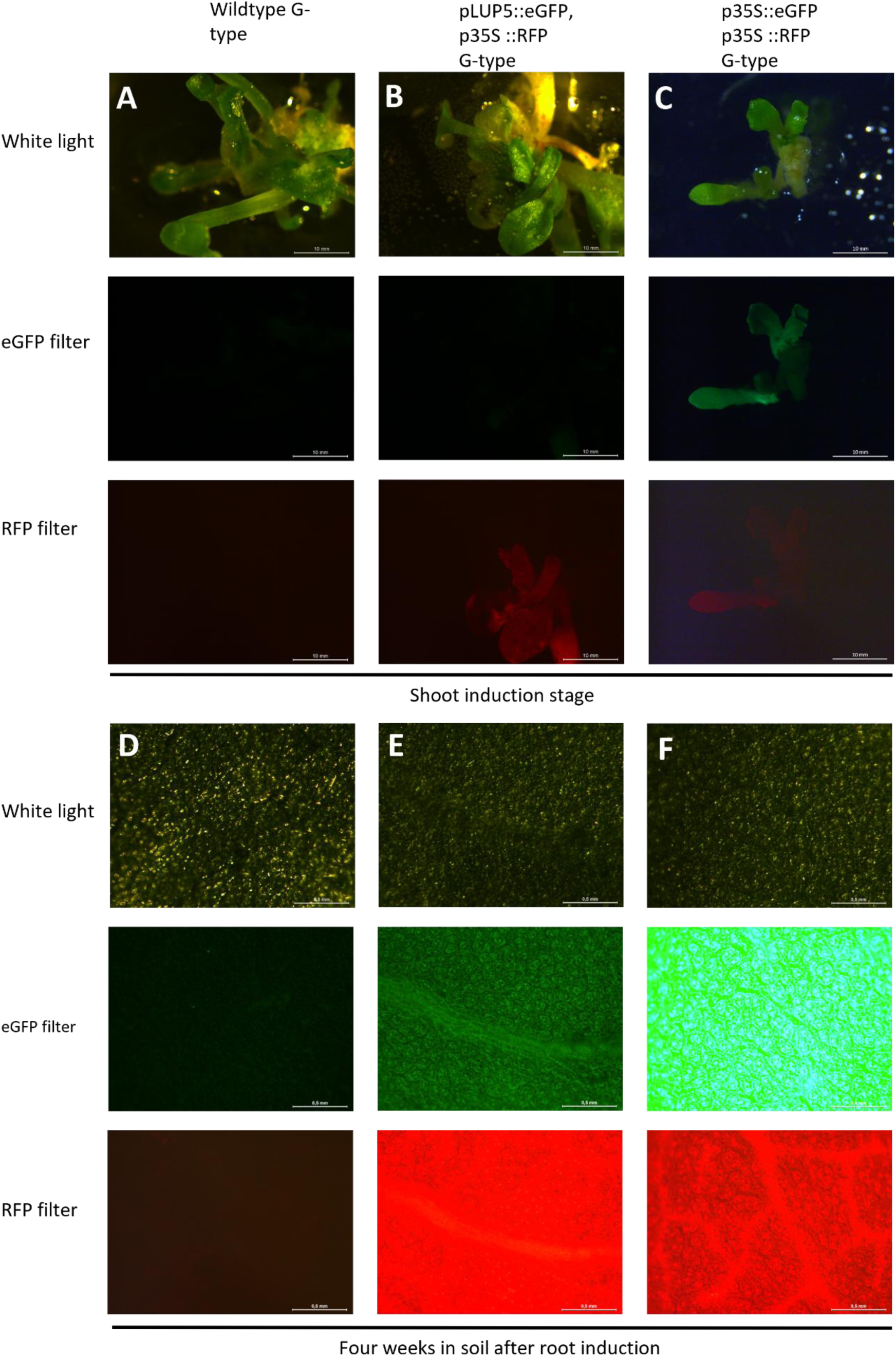
Native *LUP5* promoter supports higher transformation efficiency via moderate expression in *B. vulgaris*. During the shoot induction stage, no detectable eGFP fluorescence was observed in the pLUP5::eGFP lines (B). Four weeks after root induction, eGFP fluorescence could be detected in the pLUP5::GFP lines (D, E). However, GFP expression driven by the 35S promoter was still much stronger, as evident by the saturated image F, acquired with the same exposure time. RFP, driven by the 35S promoter, was used for screening transgenic plants by microscopy. A, D: Wild-type plants under white light, eGFP, and RFP filters; B, E: pLUP5::eGFP transformed G-type plants under white light, eGFP, and RFP filters; C, F: p35S::eGFP transformed G-type plants under white light, eGFP, and RFP filters; Scale bars: A–C, 10 mm, D–F, 0.5 mm.

Introduction of G-type *LUP5* in G-type increased a monodesmosidic hederagenin (with three hexoses) and a oleanolic acid monoglucoside (with two hexoses) in both p35S::LUP5 and pLUP5::LUP5 transformed G-type plants, retention times 8.5, and 11.0 min, respectively (Fig. 5, A to C). Monodesmosidic oleanolic acid (with 3 hexose, retention time at 10.7 min) increased in all pLUP5::LUP5 transformed G-type lines (Fig. 5C). Twenty-six additional saponins remained unchanged (Supplementary Table S8), and the three slightly elevated saponins remained at low abundance relative to other saponins (Fig. 5A-C, Supplementary Table S8).

**Fig. 5.**
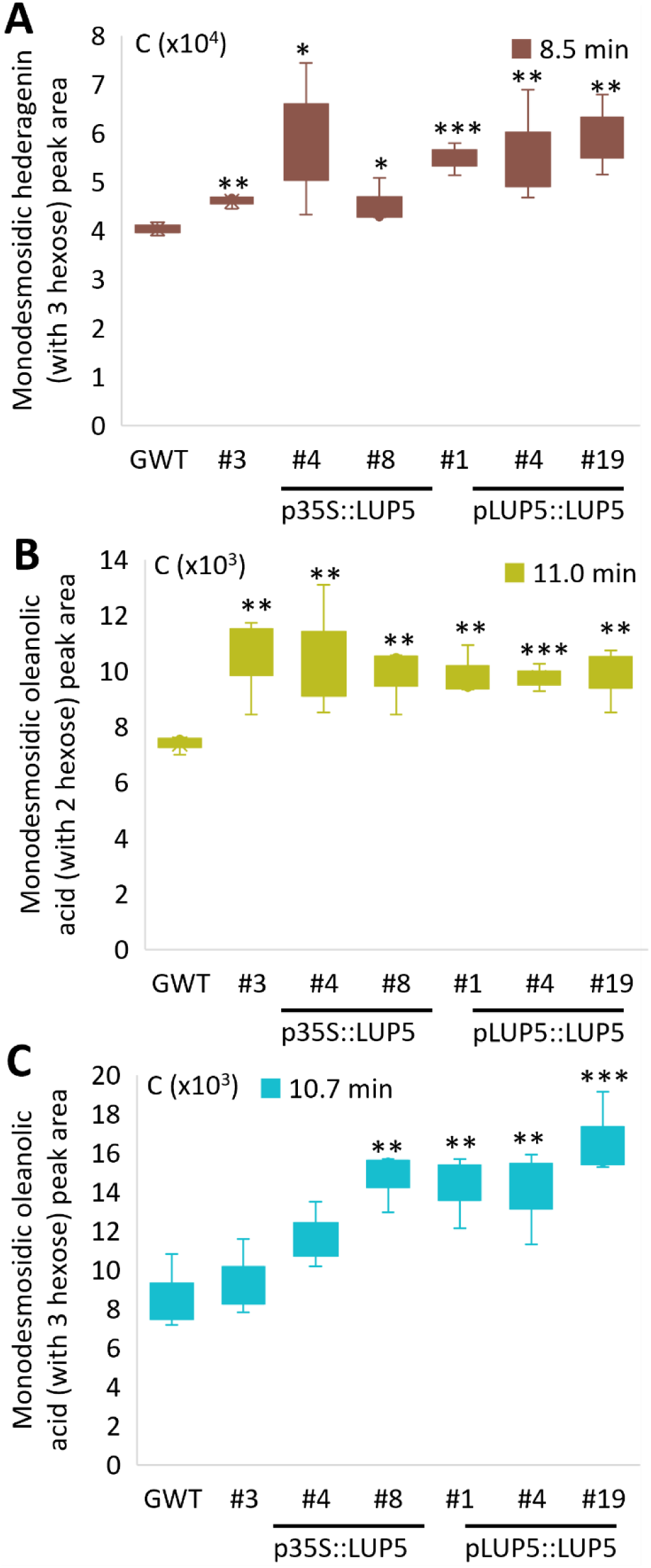
Expression of G-type *LUP5* increased glycosylated oleanolic acid and hederagenin levels in G-type *B. vulgaris*. A) Monodesmosidic hederagenin (with 3 hexose) peak area (EIC: *m/z* = 1003.5115 ± 0.2). B) Monodesmosidic oleanolic acid (with two hexoses) peak area (EIC: *m/z* = 617.4059 ± 0.2). C) Monodesmosidic oleanolic acid (with 3 hexose) peak area (EIC: *m/z* = 987.517 ± 0.2). GWT, wildtype G-type *B. vulgaris*; p35S::LUP5, p35S::LUP5 transformed G-type *B. vulgaris*; pLUP5::LUP5, pLUP5::LUP5 transformed G-type *B. vulgaris*. Error bars represent the standard deviation of the mean from three individual plants. Statistical significance was assessed using Student’s *t*-test: *(p<0.5), ** (p < 0.05) and *** (p < 0.005).

In conclusion, stable transformation of G-type *LUP5* in P-type significantly increased one oleanolic acid monoglucoside (with one hexose) and three hederagenin-derived monodesmosidic saponins (each with two hexoses) by approximately 50% to fourfold, depending on the promoter and compound (Fig. 3, A and B). Notably, the oleanolic acid monoglucoside, eluting at 9.5 minutes, was detected exclusively in the transformed P-type (Fig. 3B, Supplementary Table S5). The most abundant hederagenin-derived saponin in P-type eluted at 11.5 min, whereas hederagenin cellobioside (eluted at 12.3 min) remained the dominant saponin in G-type (Fig. 3A, Supplementary Table S6 and S8). Introduction of G-type *LUP5* in G-type slightly increased saponin levels, but their concentrations remained low compared to other oleanolic acid- and hederagenin-derived compounds (Fig. 5, A to C, Supplementary Table S8). Expression of G-type *LUP5* by its native promoter resulted in higher transformation efficiency and induced greater accumulation of β-amyrin-derived sapogenins and their glycosylated saponins, despite lower LUP5 transcript levels compared to the 35S promoter (Fig. 3, A to C, Fig. 5, A to C, Table 1, Supplementary Table S7).

### *CYP72A552* silencing decreased to 40% hederagenin derived saponin accumulation in G-type *B. vulgaris*

To evaluate the impact of G-type *CYP72A552* silencing on insect resistance, a choice insect feeding assay was conducted as described above. However, neither the transgenic G-type nor the wild-type G-type plants were consumed by diamondback moth larvae as measure after 7.5 hours (Fig. 6G). To assess the impact of *CYP72A552* downregulation on saponin and sapogenin levels in G-type *B. vulgaris*, a metabolomic analysis was performed as described above. Silencing of *CYP72A552* reduced the accumulation of six saponins in the G-type, including hederagenin monoglucoside and three other hederagenin-derived monodesmosidic saponins (each with two hexoses), monodesmosidic gypsogenin (with two hexoses), and one unknown saponin (Fig. 6, A to E). MS/MS analysis indicated that the unknown saponin contains a sapogenin of 457 m/z with one hexose and two methylpentoses (Fig. 6E). Twenty-three other saponins, including seven glycosylated oleanolic acids, remained unchanged (Supplementary Table S9). When the sapogenin levels of *CYP72A552*-silenced line #13 and wild-type G-type *B. vulgaris* were compared, the analysis revealed a 40% reduction in total hederagenin, while total and free oleanolic acid levels remained unchanged, consistent with the saponin data (Fig. 6A–F).. Lupeol and β-amyrin remained undetectable in G-type *B. vulgaris* (Supplementary Table S7). In conclusion, silencing *CYP72A552* significantly reduced hederagenin biosynthesis and the accumulation of its glycosylated derivatives in G-type *B. vulgaris*.

**Fig. 6.**
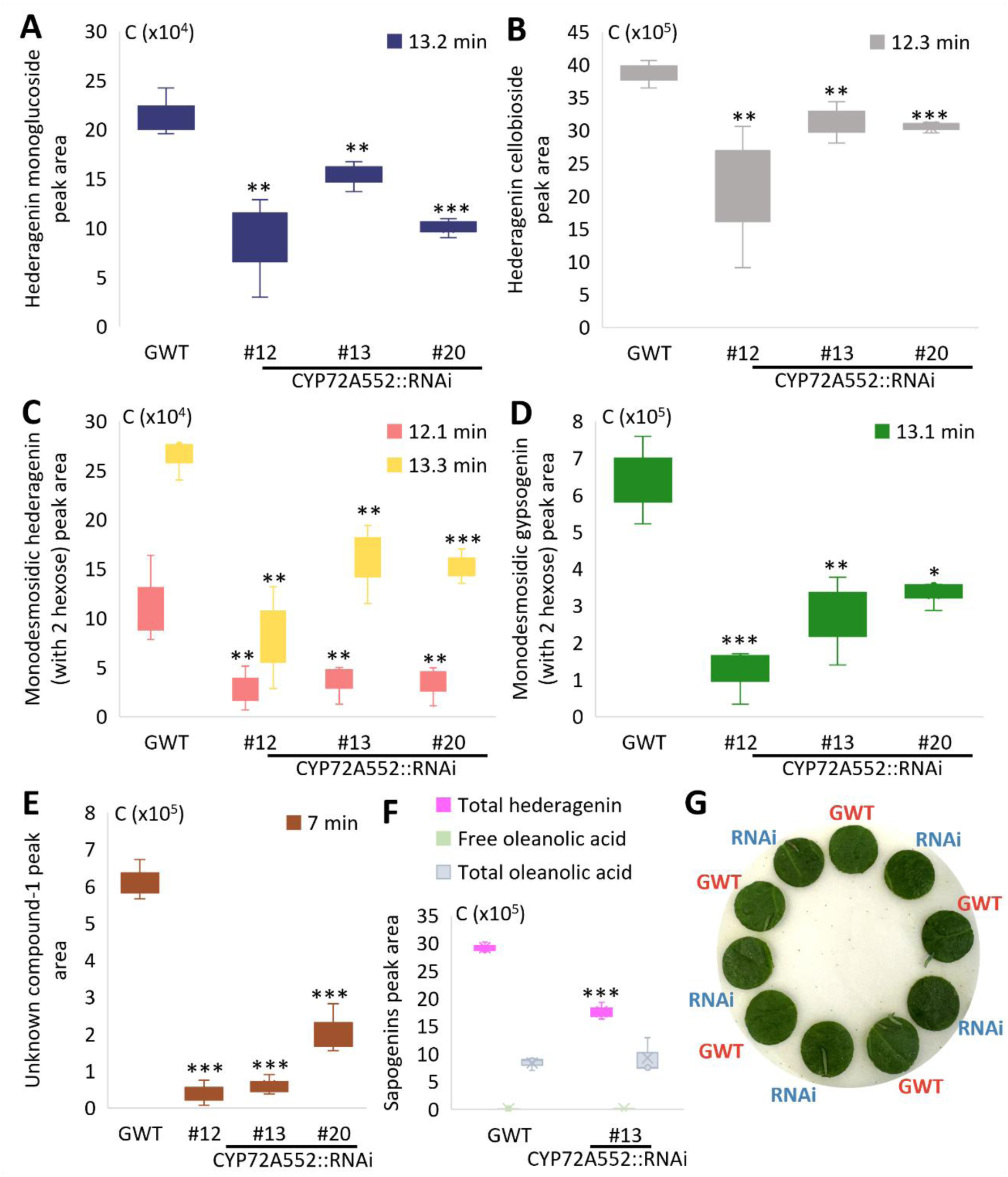
Silencing *CYP72A552* decreased hederagenin and its derived saponins in G-type *B. vulgaris*. A) Hederagenin monoglucoside peak area (EIC: *m/z* = 679.4063 ± 0.2). B) Hederagenin cellobioside peak area (EIC: *m/z* = 841.4591 ± 0.2). C) Monodesmosidic hederagenin (with two hexoses) peak area (EIC: *m/z* = 841.4591 ± 0.2). D) Monodesmosidic gypsogenin (with two hexoses) peak area (EIC: *m/z* = 839.4409 ± 0.2). E) unknown compound-1 peak area (EIC: *m/z* = 901.2429 ± 0.2). F) Peak area of sapogenins, including total hederagenin (EIC: *m/z* = 437.3425 ± 0.2), free and oleanolic acid (EIC: *m/z* = 439.3593± 0.2). G) Example of the insect choice feeding assay, and the resulting leaf condition after 7.5 hr insect exposure. GWT, wildtype P-type *B. vulgaris*; CYP72A552::RNAi, *CYP72A552* silenced G-type *B. vulgaris*. Error bars represent the standard deviation of the mean from three individual plants. Statistical significance was assessed using Student’s *t*-test: *(p<0.5), ** (p < 0.05) and *** (p < 0.005).

## Discussion

Deciphering the genetic and biochemical bases of natural insect resistance is essential for our basic understanding of chemical ecology and for addressing global challenges in crop protection and sustainability. Our study contributes to this effort by elucidating how structural diversification in plant specialized metabolites mediates ecological defense and by identifying regulatory elements that can be harnessed for future resistance breeding.

Triterpenoid saponins are key mediators of plant–insect interactions and have attracted attention due to their high chemical diversity and structure-dependent bioactivity and unique modes of action. Key factors influencing insect resistance include the structure of the saponin, the composition of the sapogenin backbone, the types of sugar moieties, the length and number of glycoside chains, and the sites of glycosylation (Gao et al., 2011; Liu et al., 2019; Tian et al., 2021). Prior metabolic engineering efforts have focused on modifying saponin composition by disrupting biosynthetic genes in legumes (Confalonieri et al., 2021; Hodgins et al., 2024). For example, Confalonieri et al. (2021) knocked out *CYP93E2*, which hydroxylates β- amyrin at the C-24 position, in *Medicago truncatula*, resulting in transgenic lines that no longer produced non-hemolytic soyasapogenols saponins but instead redirecting metabolic flux toward the synthesis of hemolytic saponins. Hodgins et al. (2024) knocked out β-amyrin synthase *BAS1* in pea (*Pisum sativum*), leading to a 99.8% reduction in saponin content in the seeds. However, these studies did not address how targeted saponin engineering influences ecological traits such as herbivore resistance or consumption.

We have previously transiently expressed saponin biosynthetic genes in *Nicotiana benthamiana* (Khakimov et al., 2015; Liu et al., 2019). An unexpected drawback of this transient tobacco system was that hederagenin was further metabolized by endogenous tobacco enzymes, which led to the accumulation of hederagenin-3-*O*-monoglucoside to levels far below those naturally occurring in *Barbarea vulgaris* (Liu et al., 2019). When oleanolic acid monoglucoside was transiently produced in *N. benthamiana*, mainly non-toxic bidesmosidic saponins accumulated, suggesting that the toxic monoglucosides were further glycosylated and thereby detoxified by the plant (Khakimov et al., 2015). In addition, the diamondback moth is a crucifer specialist and will not feed on *N. benthamiana*. To overcome these major obstacles, we metabolically engineered the saponin biosynthetic pathway directly in the wild crucifer *B. vulgaris* to test whether specific triterpenoid profiles confer resistance to insect herbivory.

We hypothesized that differences in *LUP5* expression between G- and P-type *B. vulgaris* underlie their distinct saponin profiles and insect resistance, based on quantitative trait locus (QTL) analysis and insect feeding assays with saponin-treated leaf disks (Augustin et al., 2012; Khakimov et al., 2015; Nielsen et al., 2010; Kuzina et al., 2011). Our metabolic engineering strategy demonstrates that the susceptibility of the P-type to diamondback moth feeding is primarily due to its lack of *LUP5* expression. LUP5 encodes an oxidosqualene cyclase that produces β-amyrin, the precursor of oleanolic acid and hederagenin-derived saponins, both implicated in insect deterrence. Interestingly, our results further suggest that glycoside linkage exerts less influence on feeding behavior than the sapogenin backbone itself.

Variation in oxidosqualene cyclase product specificity may be a key driver of saponin-profile differences between the two *B. vulgaris* ecotypes. Insect-susceptible P-type plants predominantly produce lupeol-derived saponins, reflecting their higher expression of *LUP2*, while G-type plants express *LUP5* and accumulate β-amyrin-derived saponins, including oleanolic acid and hederagenin cellobioside, both contributing to insect resistance (Augustin et al., 2012; Khakimov et al., 2015; Liu et al., 2019; Günther et al., 2021). By combining gene transfer with ecological assays, our study provides a framework for linking specialized metabolism with ecological function in a non-model plant system.

To test this hypothesis in an *in planta* system, the β-amyrin synthase *LUP5* was expressed in both the insect resistant G-type and insect susceptible P-type *B. vulgaris*. In the otherwise insect susceptible P-type, this resulted in up to a 95% reduction in leaf consumption and increased levels of three hederagenin-derived monodesmosidic saponins (each with two hexoses) and one oleanolic acid monoglucoside (Fig. 2A, Fig. 3, A and B). In wildtype *B. vulgaris* G-type, hederagenin cellobioside is the predominant saponin, while only small amounts of the other two hederagenin-derived saponins were detected at retention times of 10.7 and 11.5 min (Fig. 5, A and B, Supplementary Table S8). In contrast, in the *LUP5*-transformed P-type, the monodesmosidic hederagenin eluting at 11.5 min is more abundant than the other two (Fig. 3A). This variation highlights differences in saponin glycosylation patterns between P- and G-types, suggesting that these three hederagenin-derived monodesmosidic saponins, particularly the one detected at 11.5 min, may contribute to insect resistance in the transformed P-type plants. Hederagenin 3-*O*-glucoside exhibits seven-fold stronger feeding reduction to diamondback moth and tobacco hornworm compared to oleanolic acid 3-*O*-glucoside (Liu et al., 2019). This enhanced efficacy has been linked to C-23 hydroxylation in hederagenin, which causes the C-3 glucose moiety to adopt a different orientation relative to the sapogenin backbone compared to oleanolic acid (Cárdenas, Almeida and Bak, 2019; Liu et al., 2019). Considering the bioactivity and the relatively low increase of oleanolic acid monoglucoside compared to hederagenin-derived monodesmosidic saponins in *LUP5*- transformed P-type suggest that there is a CYP72A552 ortholog in the P-type that effectively transforms the oleanolic acid to hederagenin. This hederagenin is then glycosylated by different UGTs than in the G-type *B. vulgaris* to other linkage types than the 1→4 linkage cellobiosides that predominantly accumulate in the G-type.

Based on our findings, *LUP5* is identified as the key determinant for insect resistance differences between G- and P-type *B. vulgaris*. Furthermore, three hederagenin-derived monodesmosidic saponins (each with two hexoses) may play a role in insect resistance in *LUP5*-transformed P-type, particularly the most abundant saponin at retention time 11.5 min. These findings offer new mechanistic insight into plant–insect coevolution and establish a model for dissecting genotype–phenotype relationships in plant chemical defense. Future work comparing the insect resistance differences between the elucidated structures of these three hederagenin-derived monodesmosidic saponins could provide further insights into how glycosylation patterns affect saponin bioactivity. Glycoside linkage impacts cytotoxicity of saponins in *in vivo* assays (Chwalek et al., 2006). However, how glycoside linkage affects insect resistance remains unknown. Unfortunately, due to their low concentration, isolating and purifying these compounds for bioactivity testing remains challenging. Future research should focus on identifying candidate glycosyltransferases that produces a variety of glycoside linkage with the aim to produce defense-related hederagenin-derived monodesmosidic saponins *in planta*.

Interestingly, when *LUP5* expression was driven by the constitutive 35S promoter, *LUP5* was expressed at approximately sixteen-fold higher levels than in the best *pLUP5::LUP5* transgenic line and also exceeded expression levels in both wild-type P- and G-type plants (Fig. 2C). However, this high expression did not translate into significantly increased saponin production (Fig. 2, A and B, Fig. 5, A to C). In contrast, using the native *LUP5* promoter not only resulted in approximately two-fold higher transformation efficiency (Table 1) but also led to greater accumulation of hederagenin and its glycosides in the P-type background despite the overall much lower expression profile (Fig. 3, A and C). These findings indicate that expression strength alone does not determine metabolic output.

We propose that high and premature expression of *LUP5* under the 35S promoter leads to early toxic β-amyrin accumulation, thereby reducing transformation efficiency (Fig.2 and Fig. 3). In *B. vulgaris*, *LUP5* is expressed most strongly in leaves compared with roots and petioles (Khakimov et al., 2015). To better understand the observed differences in transformation efficiency, saponin and sapogenin accumulation, and *LUP5* expression levels between the native and constitutive promoters, we compared promoter activity and eGFP fluorescence intensity during plant regeneration and early growth. Microscopy observations support this interpretation: the 35S promoter was already active during shoot induction, whereas the native *LUP5* promoter became detectable only four weeks after transfer to soil following root induction (Fig. 2). Such premature activation likely causes β-amyrin to accumulate before downstream pathway genes are expressed, disturbing metabolic balance through substrate depletion or feedback inhibition. By contrast, delayed activation under the native *LUP5* promoter allows coordinated expression with downstream enzymes, promoting efficient accumulation of of saponins that are non-toxic to the plant.

Thus, temporal coordination of gene expression emerges as a key determinant of successful metabolic engineering. Although constitutive promoters such as 35S are widely used to achieve strong expression, their untimed activity can impose metabolic burdens and reduce transformation efficiency. Similar effects have been reported in other plant systems: in cotton (*Gossypium hirsutum*), continuous 35S activity throughout development caused unintended metabolic load (Sunilkumar et al., 2002), while in maize (*Zea mays*), the *ZmUbi1* promoter achieved higher transformation efficiency and better tissue-culture survival than 35S despite lower transcript levels (Beringer et al., 2017). These findings reinforce that expression timing, rather than transcriptional magnitude, is crucial for maintaining metabolic homeostasis and efficient pathway flux.

Taken together, although the 35S promoter can drive high levels of transgene expression, its continuous activity often disrupts metabolic balance, decreases transformation efficiency, and induces pleiotropic effects on plant development. In contrast, the native LUP5 promoter confines expression to appropriate developmental stages, providing precise metabolic control and coordinated biosynthetic regulation. These results highlight that promoter choice governs metabolic outcomes not merely through expression strength but through temporal alignment with downstream biosynthetic processes, thereby ensuring efficient *in planta* production of defense-related hederagenin-derived monodesmosidic saponins.

We had anticipated that silencing of *CYP72A552* in G-type would reduce insect consumption as hederagenin derived saponins are more bioactive than those derived from oleanolic acid (Liu et al., 2019) . However, our results show that while silencing of *CYP72A552* decreased hederagenin biosynthesis by up to 40%, leading to a significant reduction in hederagenin- and gypsogenin (which is hederagening that has been further oxidized at C-23 to the ketone)-derived monodesmosidic saponins. Specifically, the bioactive hederagenin monoglucoside, hederagenin cellobioside and monodesmosidic gypsogenin (with two hexoses) were decreased (Fig. 6, A to D). The decrease in monodesmosidic gypsogenin and unknown compound 1 may have been caused by the reduced hederagenin biosynthesis or off-target effects (Fig. 6, D and E, Supplementary Fig. S2), possibly resulting from the high nucleotide sequence identity of eight tandemly repeated *CYP72A* homologs (Liu et al., 2019). Previous studies indicate that cellobiosides of oleanolic acid, hederagenin, and gypsogenin both correlate with insect resistance in *B. vulgaris* in a concentration-dependent manner (Shinoda et al., 2002; Agerbirk et al., 2003; Kuzina et al., 2009; Liu et al., 2019). The hederagenin cellobioside concentration in G-type *B. vulgaris* leaves is ∼eleven-fold higher than the ED_50_ (50% effective dose) for *P. xylostella* (Shinoda et al., 2002; Liu et al., 2019). Thus, even with a 40% reduction, the remaining saponin levels still exceed the toxicity threshold required to deter feeding. Likewise, *LUP5*-transformed P-type plants did not reach the full resistance potential of G-type plants, likely due to incomplete accumulation of the most potent hederagenin saponins (Fig. 2, A and B, Fig. 3, A to C, Fig. 6, A to G). Overexpression of *LUP5* in G-type plants produced only marginal increases in minor oleanolic acid and hederagenin-derived saponins, suggesting that G-type plants may already accumulate saponin at or close the acceptable threshold for accumulation (Fig. 6, A to C, Supplementary Table S8).

## Conclusion

In summary, this study demonstrates that targeted metabolic engineering of triterpenoid saponins in the wild crucifer *Barbarea vulgaris* is a robust approach to dissect the ecological roles of structurally diverse saponins and study insect resistance mediated by specialized metabolism. Expression of the β-amyrin synthase *LUP5* under its native promoter was more effective than constitutive expression, leading to higher transformation efficiency and increased accumulation of insect-deterring hederagenin-derived monodesmosidic saponins. This underscores the importance of promoter choice and precise transcriptional regulation in optimizing specialized metabolite production.

Our findings establish stable metabolic engineering of saponins not only as a tool for biotechnological enhancement of resistance traits, but also as a powerful approach to understanding the role of plant defense compounds in chemical ecology. In contrast to transient expression systems, which often result in unstable metabolite accumulation and further modification by host enzymes, stable engineering enables consistent expression of pathway genes and reliable assessment of metabolite function. We identified *LUP5* as a crucial determinant of insect resistance in *Barbarea vulgaris* and demonstrated that three specific hederagenin-derived monodesmosidic saponins, each containing two hexose units, are associated with this defense. Taken together, this work introduces a novel application of metabolic engineering to unravel the ecological functions of plant metabolites and provides promising strategies for crop protection through fine-tuned manipulation of biosynthetic pathways.

## Acknowledgements

We thank Thure Hauser for providing Plutella *xylostella*, Niels Agerbirk for providing wildtype *Barbarea vulgaris* G- and P-type seeds, Barry Cohen and Hanne Volpin for their help in setting up the transformation system, and Jack Olsen and Mariela Alejandra González Ramírez for their assistance in LCMS. This work was financially supported by Novo Nordisk Foundation (EcoSap NNF20OC0060298 to SB), the China Scholarship Council (CSC, 202108330041 to JS) and Marie Sklodowska-Curie Individual Fellowship (MSCA-IF 752437 to PC).

## Author contributions

JS, PC, and SB designed the research; JS, PC, JG, SKN and SB analysed the data and wrote the paper; JS, PC and SKN performed the research; JG provided the data analysis strategy and standards.

---

**Fig. S1.**
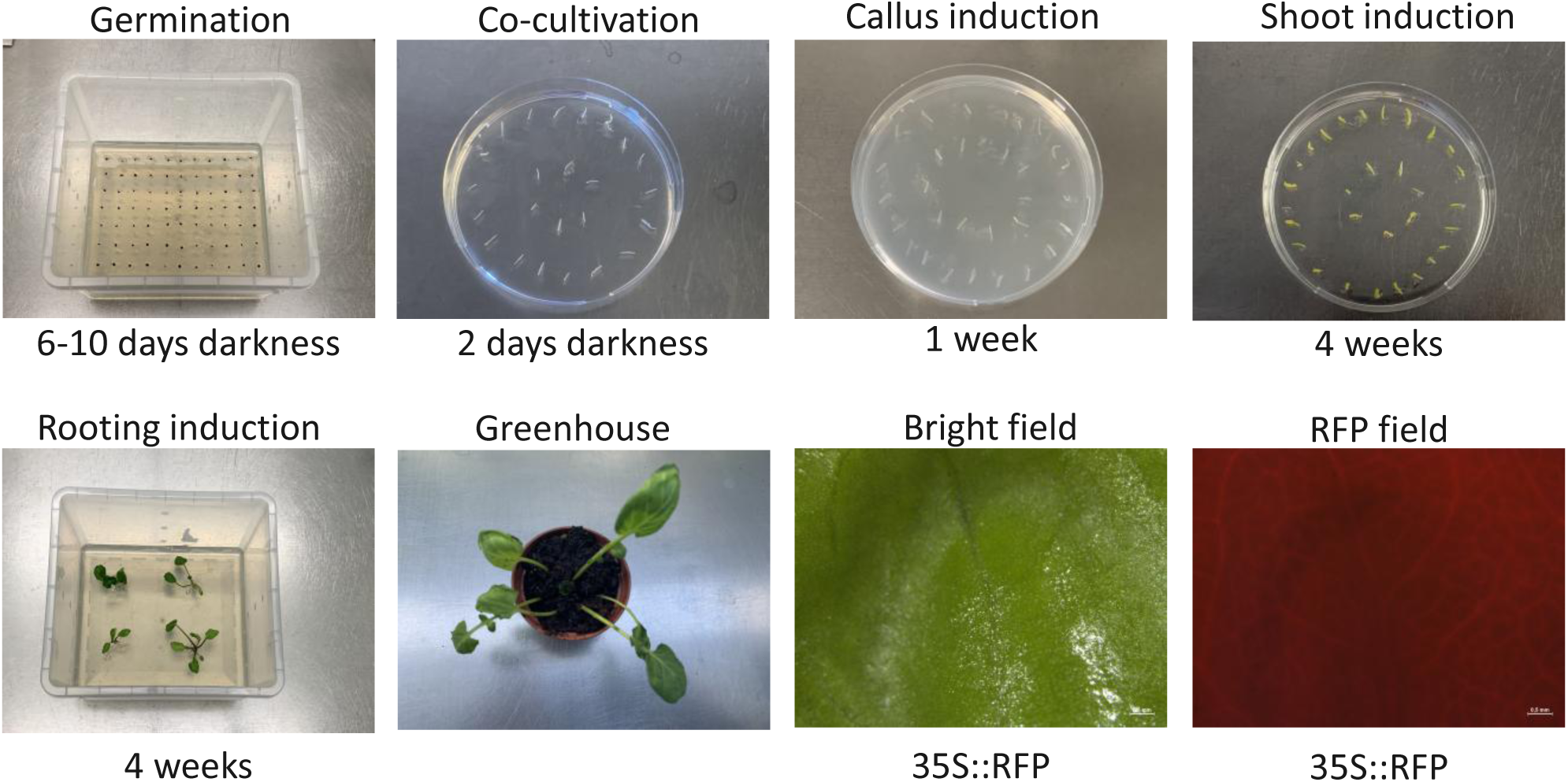
*Agrobacterium*-mediated stable plant transformation process.

**Fig. S2.**
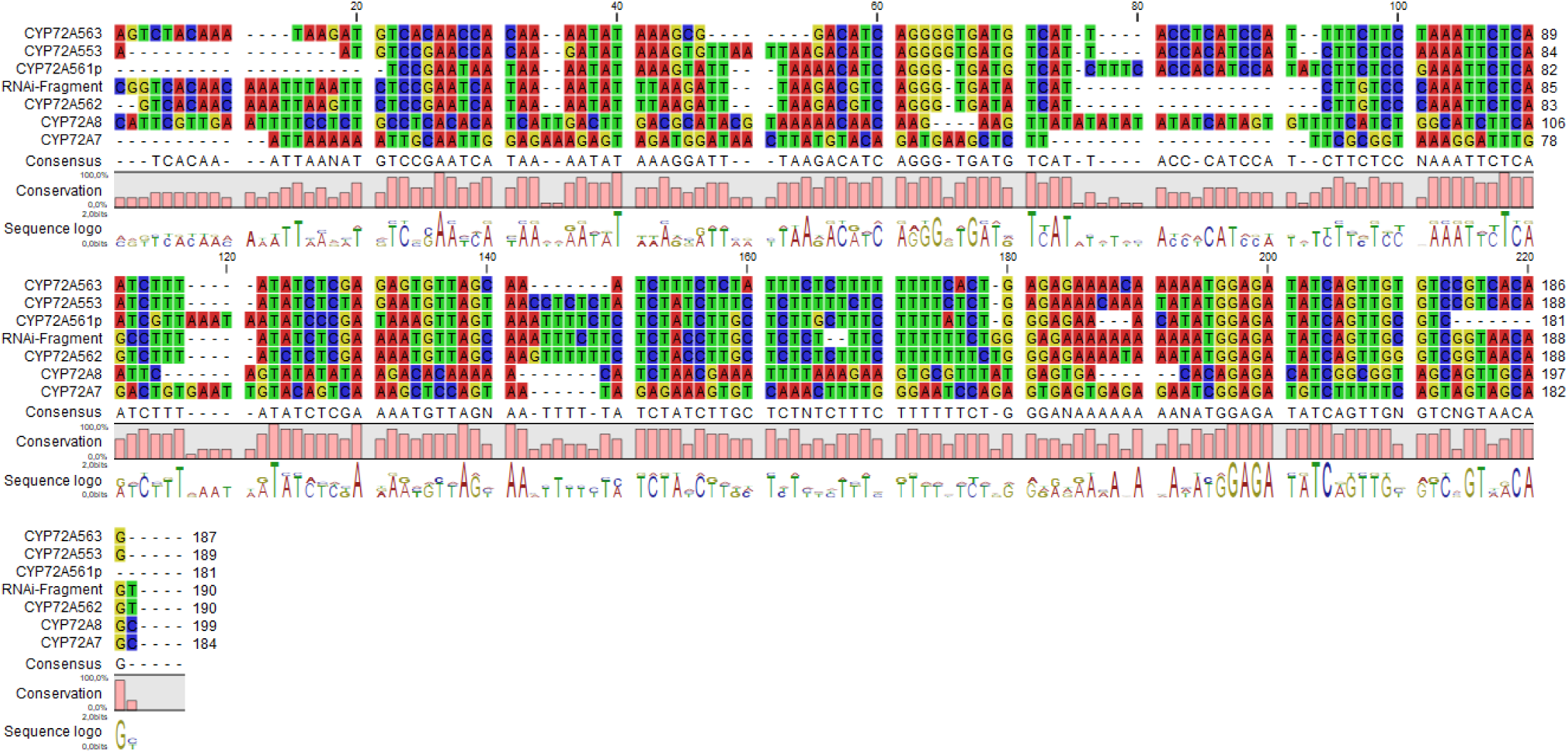
Nucleotide sequence alignment between the RNAi fragment and eight tandemly repeated *CYP72A* genes in *B. vulgaris*. Accession numbers of the eight CYP72A genes range from MH252567 to MH252574.

**Tab. S1.**
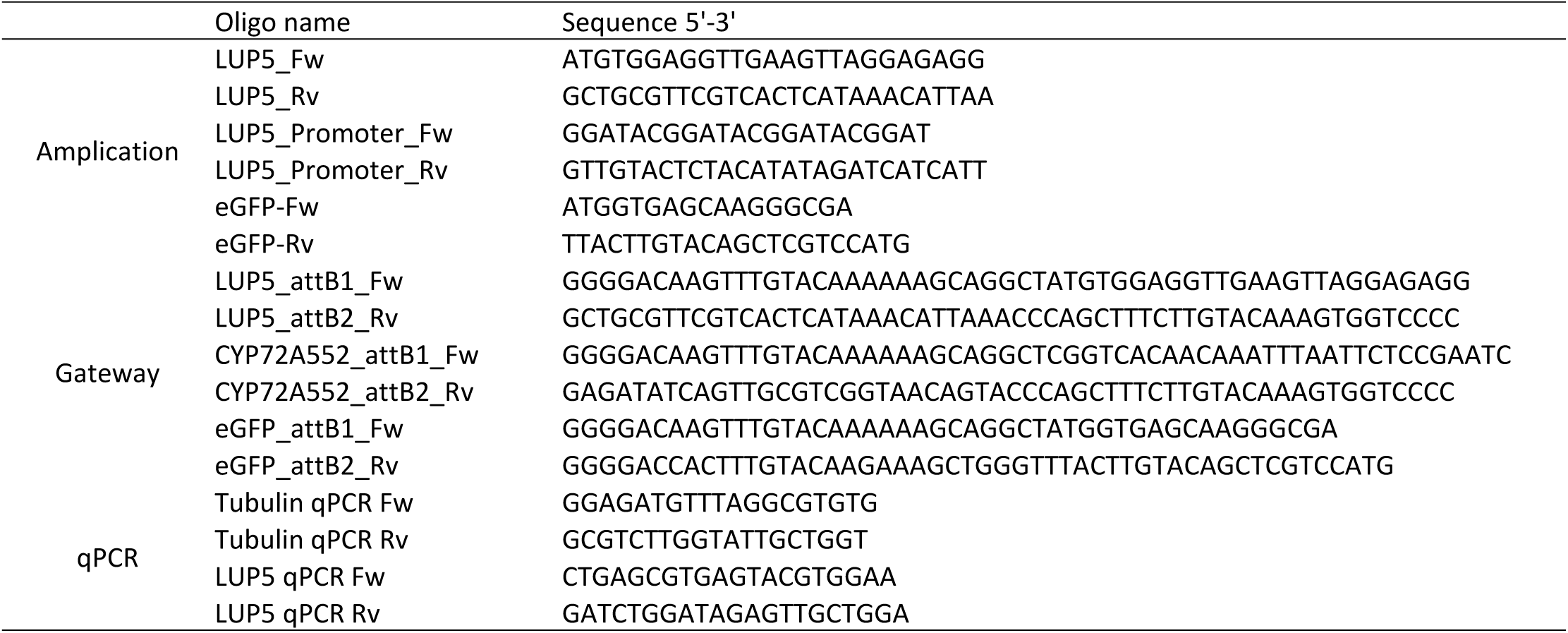
Primers used in this study.

**Tab. S2.**
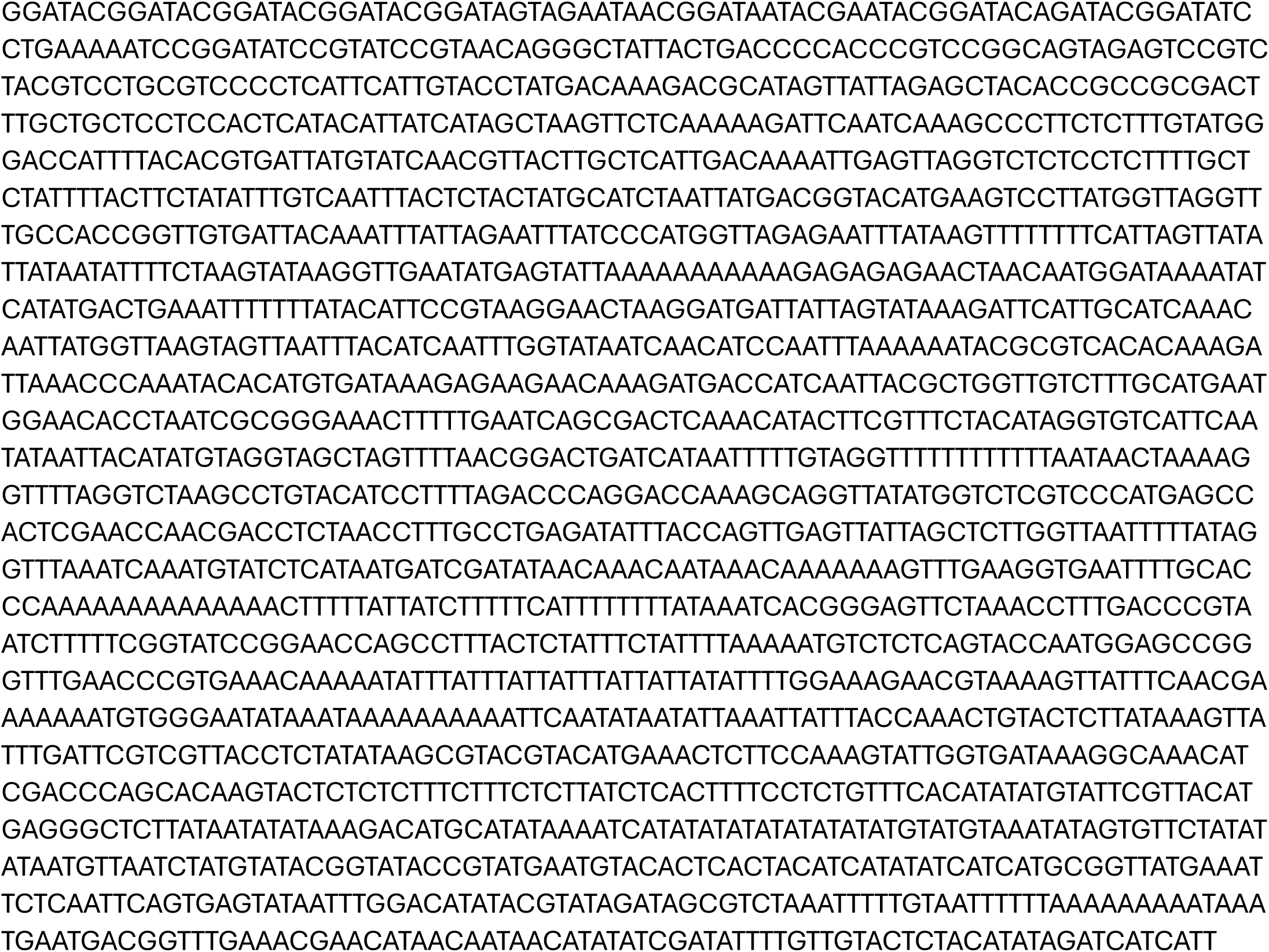
Nucleotide sequence of the G-type LUP5 promoter.

**Tab. S3.**
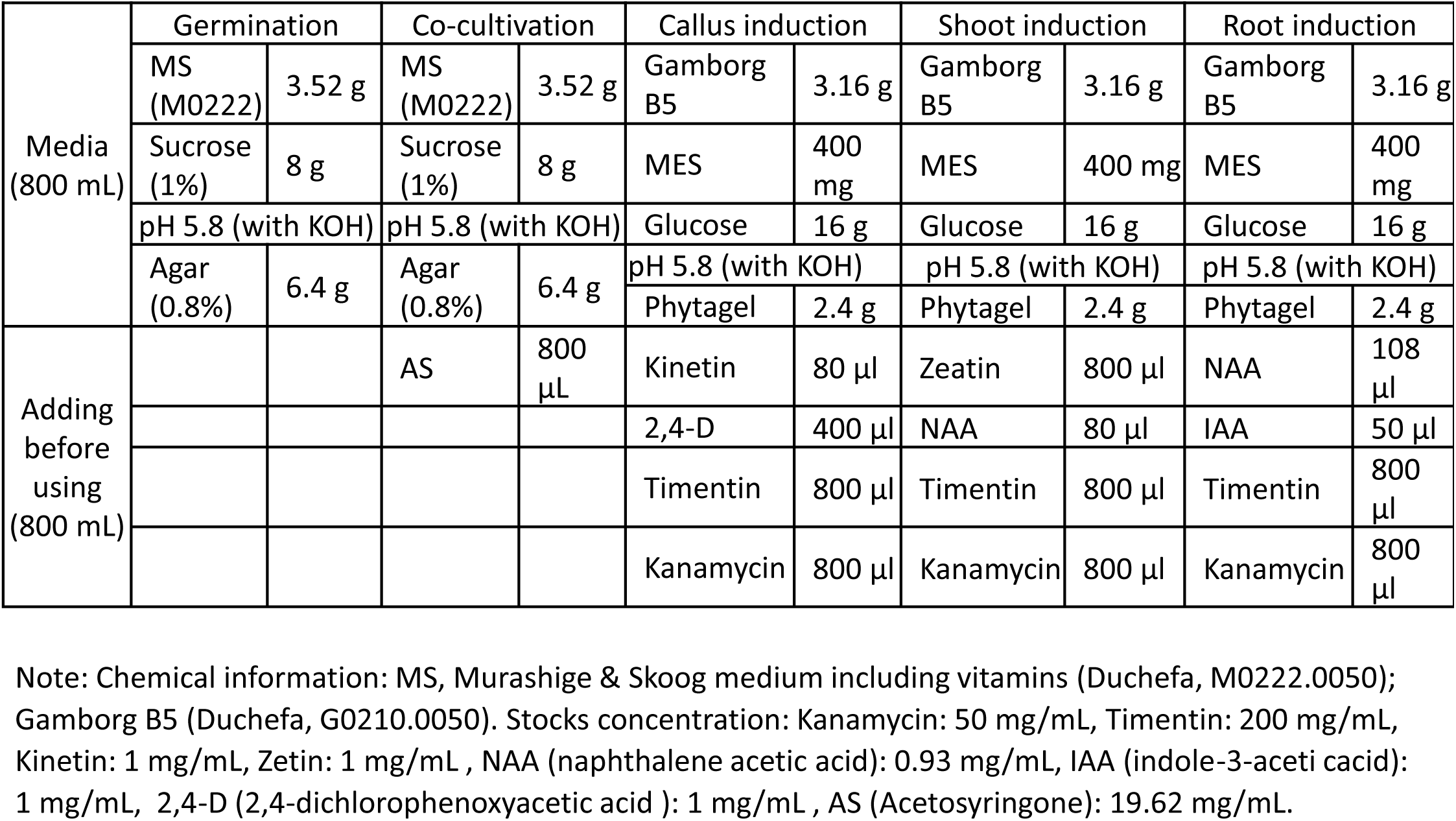
Meida for tissue culture.

**Tab. S4.** Saponins detected in P-type *B. vulgaris*. EIC, extract ion chromatogram.

| Sapog<br>enin | Saponin fragment | Sugar moieties | Formic<br>acid | Fragmentation pattern in<br>Mass Spectra | EIC±0.2 | Retention time<br>(min) |  |  |
| --- | --- | --- | --- | --- | --- | --- | --- | --- |
| 457 | 457,619,781,943, 989 | 3 x hexose | Yes | [M<sub>989</sub>-46-162-162-162=457] | 989.5404 | 8.6 | 8.9 |  |
| 447 | 447,609,755,901 | 1 x hexose; 2 x methylpentose | No | [M<sub>901</sub>-146-146-162=447] | 901.2429 | 6.3 |  |  |
| 473 | 473,635,797,959, 1005 | 3 x hexose | Yes | [M<sub>1005</sub>-46-162-162-162=473] | 1005.5337 | 8.1 | 8.4 |  |
| 473 | 473,635,797,843 | 2 x hexose | Yes | [M<sub>843</sub>-46-162-162=473] | 843.4798 | 8.6 |  |  |
| 471 | 471,633,795,841 | 2 x hexose | Yes | [M<sub>841</sub>-46-162-162=471] | 841.4591 | 10.7 | 11.5 | 12.3 |
| 455 | 455,617 | 1 x hexose | No | [M<sub>617</sub>-162=455] | 617.4059 | 9.5 |  |  |

**Tab. S5.**
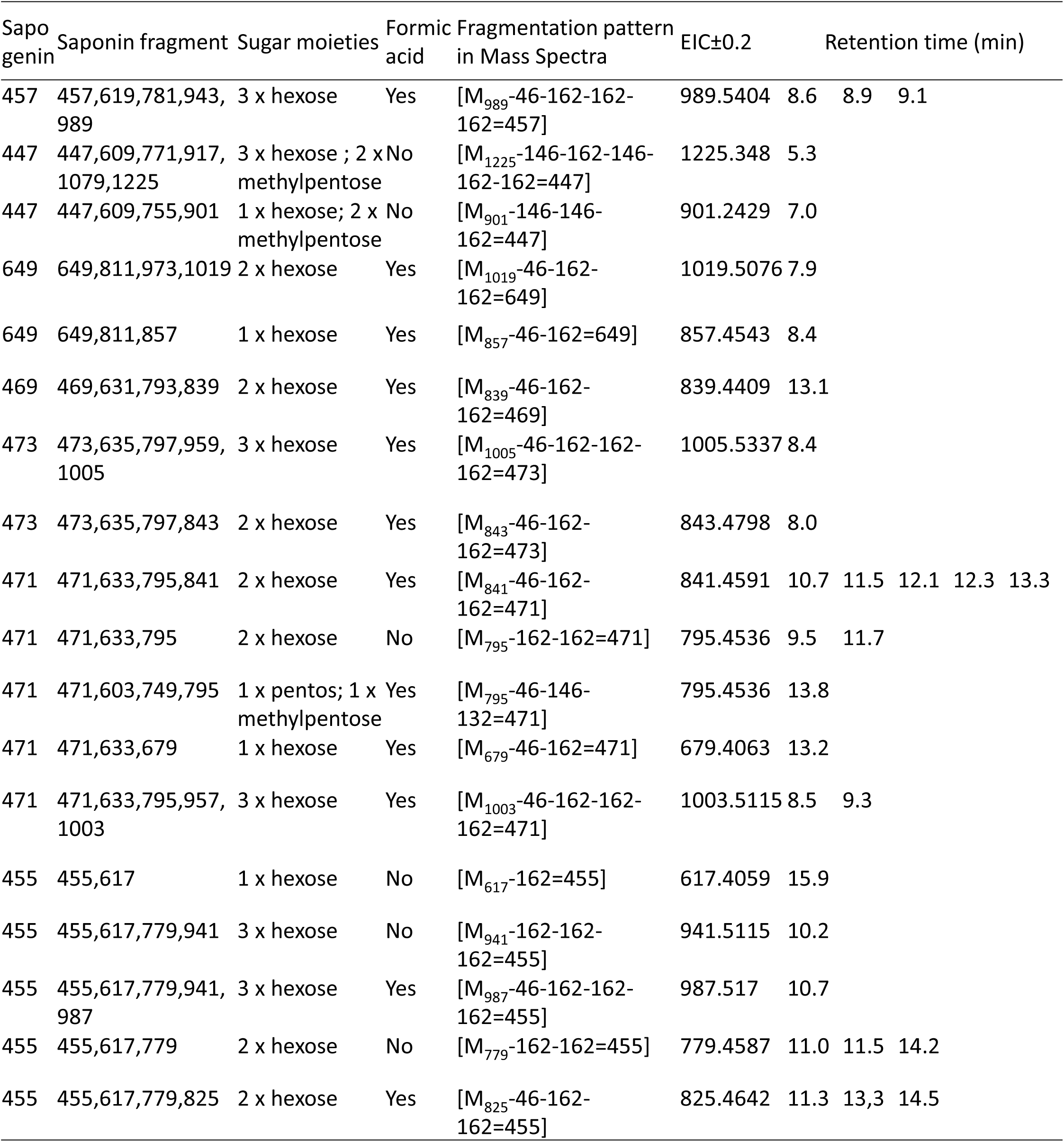
Saponins detected in G-type *B. vulgaris*. EIC, extract ion chromatogram.

**Tab. S6.**
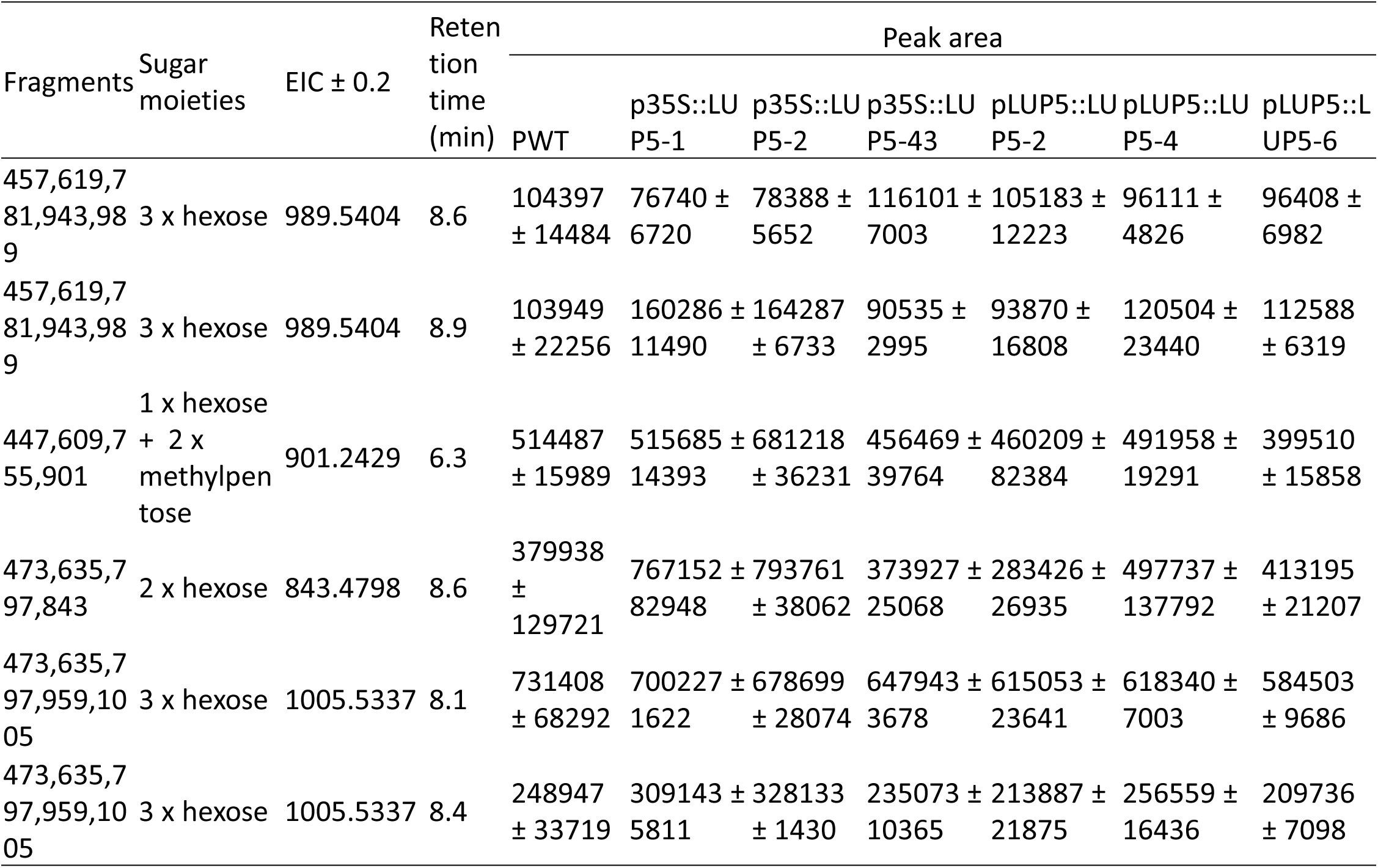
Non significantly peak area changed saponins in LUP5 transformed P-type *B. vulgaris* compared to wildtype P-type. EIC, extract ion chromatogram; PWT, wildtype P-type *B. vulgaris*; p35S::LUP5, p35S::LUP5 transformed P-type *B. vulgaris*.

**Tab. S7.** Identification and relative quantification of total and free sapogenins in *B. vulgaris*. EIC, extract ion chromatogram.

| Sapogenins | EIC±0.2 | Retention<br>time (min) | PWT | p35S::LUP5<br>-2 | pLUP5::LUP<br>5-4 | GWT | CYP72A552::R<br>NAi-13 |
| --- | --- | --- | --- | --- | --- | --- | --- |
| Total lupeol | 409.3851 | 31.1 | N/A | N/A | N/A | N/A | N/A |
| Free lupeol | 409.3851 | 31.1 | N/A | N/A | N/A | N/A | N/A |
| Total β-amyrin | 409.3851 | 32.5 | N/A | N/A | N/A | N/A | N/A |
| Free β-amyrin | 409.3851 | 32.5 | N/A | N/A | N/A | N/A | N/A |
| Total oleanolic acid | 439.3593 | 21.5 | 20884 ±<br>3963 | 45589 ±<br>6970 | 53763 ±<br>5506 | 837805 ±<br>94973 | 930214 ±<br>259536 |
| Free oleanolic acid | 439.3593 | 21.5 | N/A | N/A | N/A | 7694 ±<br>2187 | 12102 ± 3978 |
| Total hederagenin | 437.3425 | 16.7 | 37387 ±<br>13280 | 119157 ±<br>28689 | 316704 ±<br>75506 | 2923104 ±<br>69572 | 1763484 ±<br>124066 |
| Free hederagenin | 437.3425 | 16.7 | N/A | N/A | N/A | N/A | N/A |

**Tab. S8.**
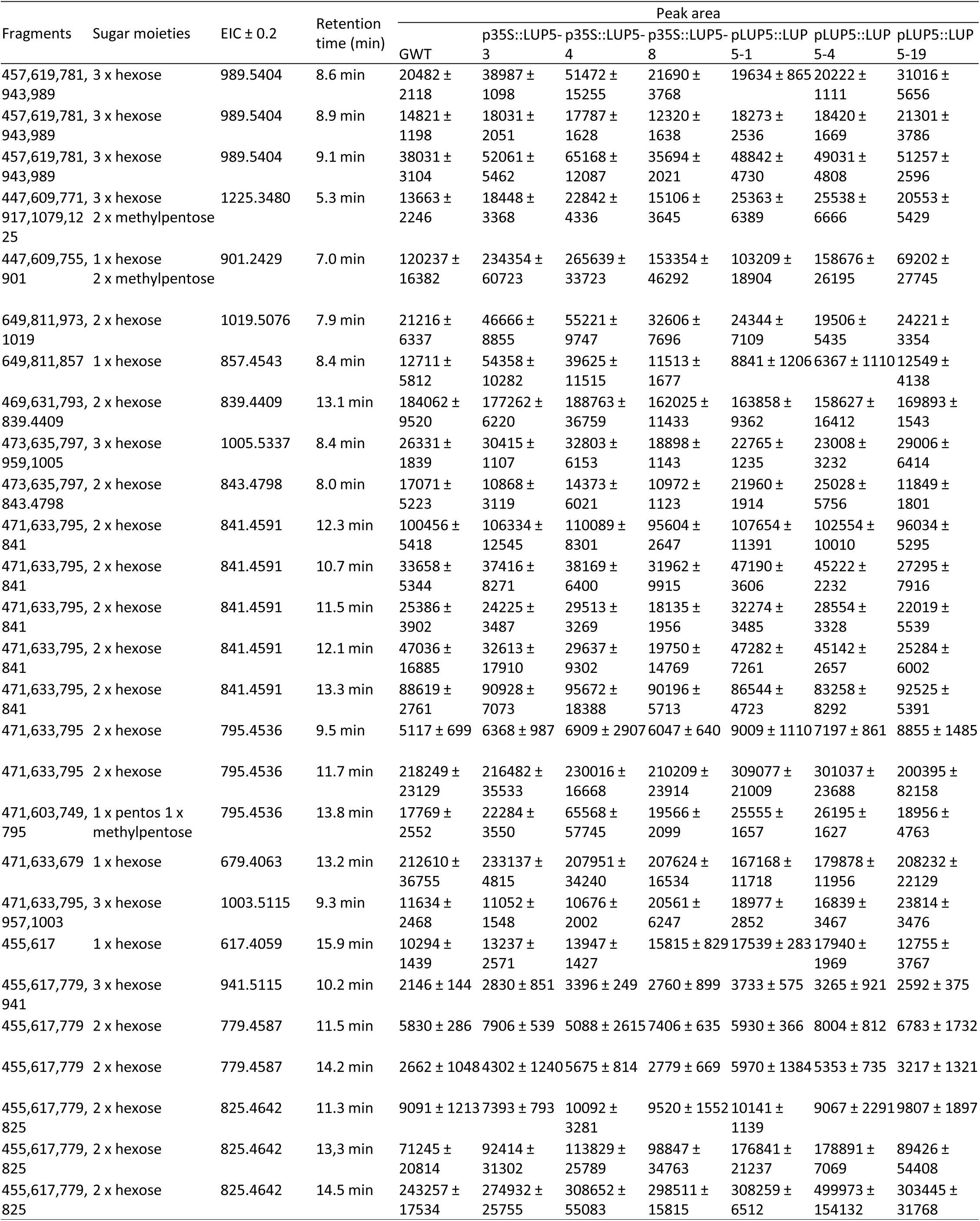
Non significantly peak area changed saponins in LUP5 transformed G-type *B. vulgaris* compared to wildtype G-type. EIC, extract ion chromatogram; GWT, wildtype G-type *B. vulgaris*; p35S::LUP5, p35S::LUP5 transformed G-type *B. vulgaris*. Hederagenin cellbioside (EIC: m/z 841.4591± 0.2, retention time 12.3 min) was quantified by after diluting sample 60 times

**Tab. S9.**
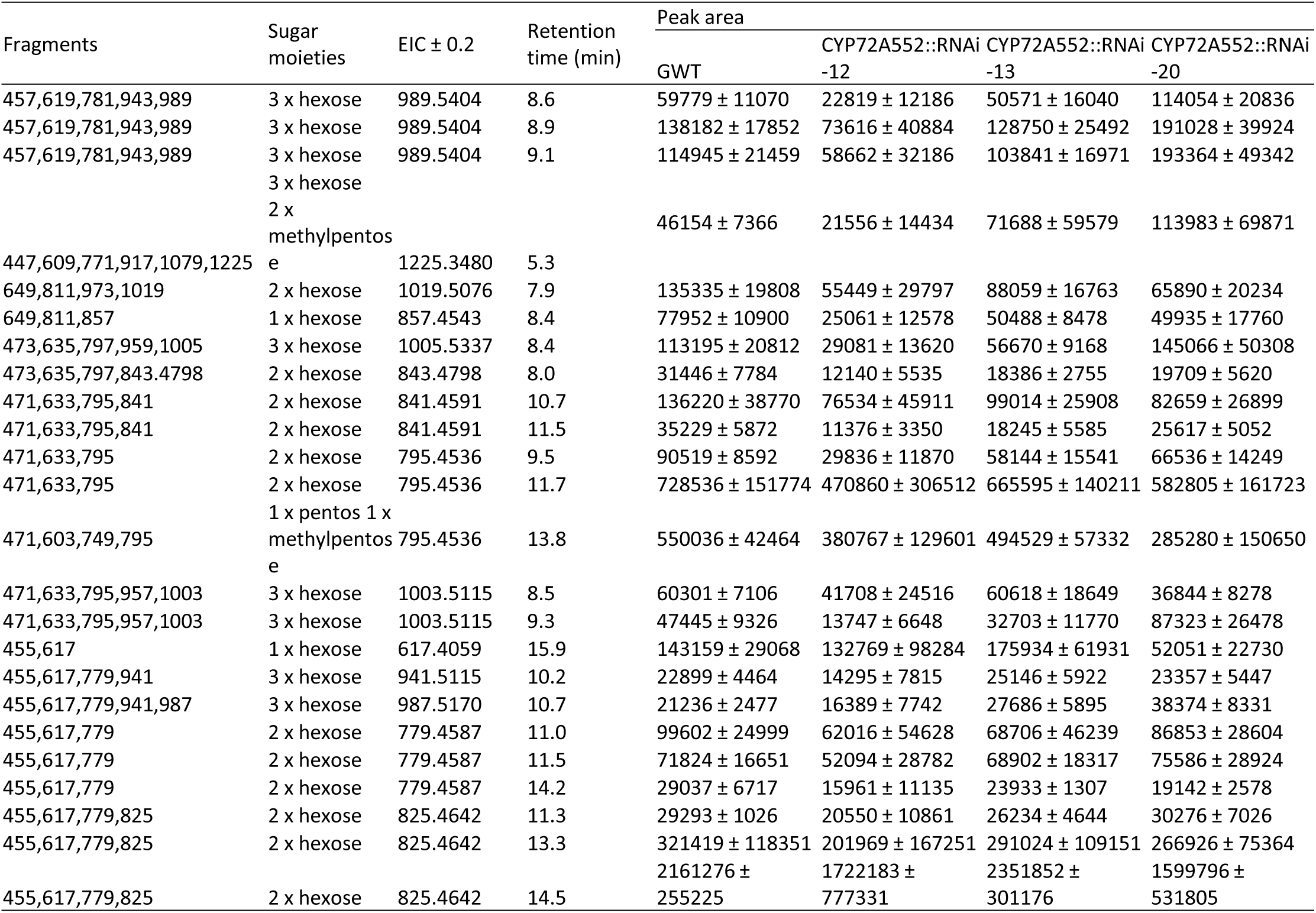
Non significantly peak area changed saponins in CYP72A552 RNAi G-type *B. vulgaris* compared to wildtype G-type. EIC, extract ion chromatogram; GWT, wildtype P-type *B. vulgaris*; CYP72A552::RNAi, CYP72A552 silenced G-type *B. vulgaris*.

